# Efflux pumps in *Chromobacterium* species and their involvement in antibiotic tolerance and survival in a co-culture competition model

**DOI:** 10.1101/562140

**Authors:** Saida Benomar, Kara C Evans, Robert L Unckless, Josephine R Chandler

**Author notes:** To whom correspondence should be addressed: Josephine R. Chandler, 1200 Sunnyside Ave. Lawrence, KS 66045.

## Abstract

Members of the *Chromobacterium* genus includes opportunistic but often-fatal pathogens and soil saprophytes with highly versatile metabolic capabilities. In previous studies of *Chromobacterium subtsugae* (formerly *C. violaceum*) strain CV017, we identified a resistance nodulation division (RND)-family efflux pump (CdeAB-OprM) that confers resistance to several antibiotics including the bactobolin antibiotic produced by the soil saprophyte *Burkholderia thailandensis*. Here, we show the *cdeAB-oprM* genes increase *C. subtsugae* survival in a laboratory competition model with *B. thailandensis.* We also demonstrate that adding sublethal bactobolin concentrations to the co-culture increases *C. subtsugae* survival, but this effect is not through CdeAB-OprM. Instead, the increased survival requires a second, previously unreported pump we call CseAB-OprN. We show the CseAB-OprN genes are transcriptionally induced in cells exposed to sublethal bactobolin concentrations and that this causes an increase in bactobolin tolerance. Induction of this pump is through a bactobolin-responsive regulator, CseR. We also demonstrate that CseAB-OprN is highly specific and sensitive to bactobolin, while CdeAB-OprM appears to have a broader range of specificities. We examine the distribution of CseAB-OprN and CdeAB-OprM in members of the *Chromobacterium* genus and find that CseAB-OprN is limited to the non-pathogenic *C. subtsugae* strains, whereas CdeAB-OprM is more widely distributed among members of the *Chromobacterium* genus including the occasional pathogen *C. violaceum.* Our results provide new information on the antibiotic resistance mechanisms of *Chromobacterium* species and use co-culture competition experiments to highlight the importance of efflux pumps in saprophytic bacteria.

**IMPORTANCE:** Antibiotic efflux pumps are best known for their role in increasing antibiotic resistance of a variety of pathogens. However, the role of these pumps in saprophytic species is much less well defined. This study describes the identification and characterization of two predicted efflux pump gene clusters in members of the *Chromobacterium* genus, which is primarily comprised of non-pathogenic saprophytes but also has several members known to occasionally cause highly fatal infections in humans. One of the predicted efflux pump gene clusters is present in every member of the *Chromobacterium* genus and shown to increase resistance to a broad range of antibiotics, including those that are clinically relevant. The other gene cluster confers resistance to a limited range of antibiotics and is found only in *Chromobacterium subtsugae*, a subset of this genus that is entirely nonpathogenic. We demonstrate these pumps can be activated by their antibiotic substrates to increase antibiotic tolerance. We also use a dual-species laboratory model to demonstrate that efflux pump activation by antibiotics is important for *C. subtsugae* to survive during competition with another antibiotic-producing bacteria. These results have implications for managing antibiotic-resistant *Chromobacterium* infections and for understanding the evolution of efflux pumps outside of the host.

## INTRODUCTION

Members of the *Chromobacterium* genus are often found in water and soil environments of tropical and subtropical regions (1), and cause occasional but often fatal infections in patients (2). Many members of this genus are also potentially useful in industry and ecological applications such as bioplastics synthesis (3, 4), hydrolyzing plastic films (5), solubilizing gold (6, 7), and pest management (8, 9). The most well-known trait shared by many members of this genus is the ability to produce a purple pigment, violacein, which has a variety of biotechnologically interesting applications including as an antimicrobial (10) and anti-cancer therapeutic (11, 12). Production of violacein and other antibiotics is controlled by *N*-acylhomoserine lactone (AHL) quorum sensing (13, 14), a type of cell-cell communication distributed widely in the Proteobacteria (15). Because production of violacein can be easily observed in cell cultures, a strain of *C. subtsugae* strain CV017 (formerly *C. violaceum* CV017) is commonly used as a biosensor of AHL signals from the environment and other bacteria (13). This strain has also been used to study mechanisms of interspecies competition using a laboratory model developed with another soil saprophyte, *Burkholderia thailandensis* (16, 17).

We are interested in understanding mechanisms that are important for competition with other strains and species in polymicrobial communities, and studying these mechanisms using relatively simple laboratory models. Our *B. thailandensis-C. subtsugae* model was previously used to show how quorum sensing-controlled antibiotics can change the dynamics of competition between species (16, 17). Here, we use this model to explore mechanisms of antibiotic defense. We initially chose *B. thailandensis* and *C. subtsugae* for our model because they each grow similarly in the conditions of the experiment, however, these species can also be isolated from similar tropical soil and water environments. Also, in our laboratory conditions, both species produce antibiotics that can kill the other species, and we wished to explore the effects of these antibiotics in our earlier experiments (16). In the case of *C. subtsugae*, the involved antibiotic(s) remain unknown, and have a relatively small but significant role in the outcome of competition. In the case of *B. thailandensis* the antibiotic is bactobolin, which has a relatively large effect in the co-culture model (16). Bactobolin is a broad-spectrum antibiotic that targets the 50S ribosomal subunit to block translation (17).

Our previous studies identified a *C. subtsugae* predicted antibiotic efflux pump gene cluster that contributes to bactobolin resistance (18). We named the predicted efflux pump CdeAB-OprM. CdeAB-OprM belongs to the nodulation division (RND) efflux pump family. RND-family efflux pumps are situated within the inner membrane and involve three proteins: an inner membrane pump, an outer membrane channel, and a periplasmic adaptor protein. Together, the proteins form a tripartite efflux pump spanning both the inner and the outer membrane (19). Drug exporters belonging to the RND family play a key role in resistance to clinically relevant antibiotics in Proteobacteria (19). Resistance to antibiotics is generally due to mutation of a transcription regulator that increases production of the efflux pump and causes the cell to become more antibiotic resistant. Although the involvement of efflux pumps in antibiotic-resistant infections has been well established, much less is known about the factors that contribute to the evolution of efflux pumps outside of a host infection. The presence of efflux pumps in many saprophytic bacteria suggest these pumps are also important in other contexts, for example for managing the toxicity of endogenous antibiotics (20). Efflux pumps might also be important for defending against antibiotics produced by other species during competition.

In this study, we use our co-culture model to explore the importance of efflux pumps during interspecies competition. We demonstrate that CdeAB-OprM increases *C. subtsugae* survival during competition with bactobolin-producing *B. thailandensis.* We also demonstrate that adding a sublethal concentration of bactobolin to the co-culture experiments increases *C. subtsugae* survival in the co-culture model, suggesting that *C. subtsugae* defense mechanisms are activated by exposure to the low doses of bactobolin. However, increased survival is not dependent on CdeAB-OprM. Instead, we find a newly identified pump, CseAB-OprN, and show this pump is important for the increased survival in co-culture. We also demonstrate the involvement of CseAB-OprN in a specific response to bactobolin and that this gene cluster is activated by sublethal bactobolin concentrations through a LysR-family regulator encoded adjacent to the *cseAB-oprN* genes, CseR. We show that the *cseAB-oprN* genes are limited to the *C. subtsugae* strains, which are not known to be pathogenic, and that the genes encoding CdeAB-OprM are widely distributed in the *Chromobacterium* genus. Together, our results describe two RND-efflux pumps found in members of the *Chromobacterium* genus and demonstrate how the evolution of antibiotic response might be important during competition with other species in the soil.

## RESULTS

### Sublethal bactobolin increases *C. subtsugae* survival in co-culture

In many bacteria, exposure to sublethal antibiotic concentrations reversibly increases antibiotic resistance, often through the activation of antibiotic efflux pumps (19), to cause induction of antibiotic tolerance. We used our *B. thailandensis-C. subtsugae* co-culture model to test whether addition of a sublethal antibiotic (bactobolin) alters the outcome of competition through induction of antibiotic tolerance. We combined exponentially growing pure cultures of *B. thailandensis* and *C. subtsugae* and added sublethal concentrations of bactobolin from *B. thailandensis* filtered culture fluid at the beginning of co-culture growth. We used 1/2 the minimal bactobolin concentration that inhibits growth (1/2 MIC, which was 0.5-0.6% culture fluid in the total co-culture). With no added bactobolin, *B. thailandensis* outcompeted *C. subtsugae* in co-cultures consistent with previous results (16, 18)(Fig. 1, ‘wild type,’ filled circles). However, when sub-MIC bactobolin was supplied in the culture, *C. subtsugae* gained an advantage over *B. thailandensis* (Fig. 1 ‘wild type,’ open circles). In identical experiments with culture fluid from a *B. thailandensis* bactobolin mutant, we observed no effect on co-culture outcomes (Fig. S1). These results show that exposure to sub-MIC bactobolin increases *C. subtsugae* survival in co-culture competition with *B. thailandensis*.

**Fig. 1.**
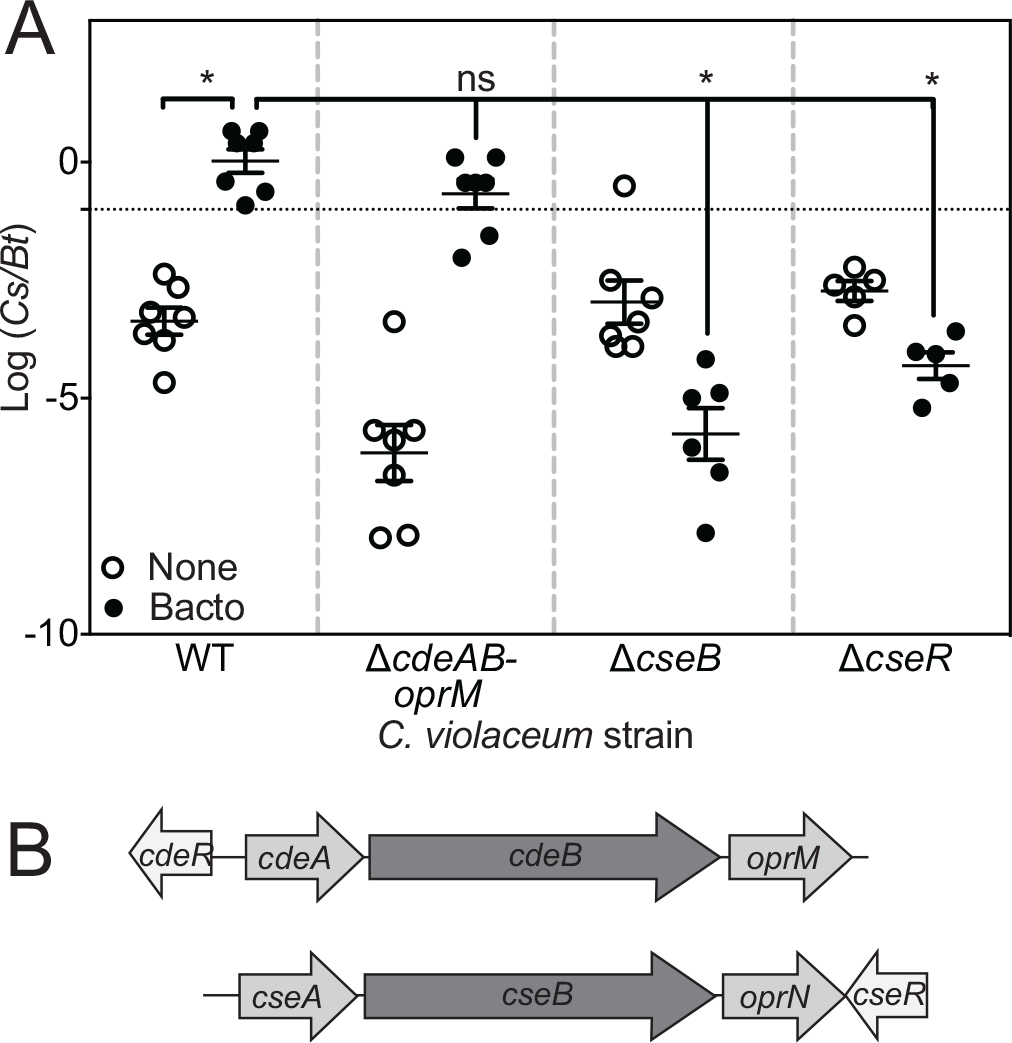
A) *B. thailandensis-C. subtsugae* competition. Co-cultures were of wild-type *B. thailandensis* (*Bt*) and *C. subtsugae* (*Cs*) CV017 or CV017 mutant strains. The black dashed line indicates the starting 1:10 ratio of *Cs* to *Bt*. The ratio of *Cs* to *Bt* (*Cs/Bt*) after 24 h was determined by selective plating and colony counts. Open circles, cultures with no added antibiotic. Filled circles, co-cultures grown with sublethal bactobolin provided by adding *B. thailandensis* filtered culture fluid (0.5-0.6%) at the start of the co-culture experiment. The solid lines represent means for each group. The vertical bars show the standard error of the mean for each group. *, statistically significant by student’s *t*-test compared with bactobolin-treated wild type (*p* < 0.001). **B) Illustration of *cdeRAB-oprM* and *cseAB-oprN-cseR* gene clusters.** Shading indicates nucleotide sequence identity. Dark grey, 80-89% identical. Light grey, 60-69% identical. The regulators (*cdeR* and *cseR*) share no identity.

### Increased survival in response to sublethal bactobolin is through a newly identified efflux pump, CseAB-OprN in *C. substugae*

Because our previous results showed CdeAB-OprM is important for bactobolin resistance, we predicted the CdeAB-OprM efflux pump might increase competitiveness in response to bactobolin (18). To test this hypothesis, we compared the competitive ability of a *C. subtsugae* Δ*cdeAB-oprM* mutant with that of wild type in our co-culture experiments. In co-cultures with no added bactobolin, we observed that the Δ*cdeAB-oprM* mutant was less competitive than wild type (Fig. 1). This result is consistent with the role of the *cdeAB-oprM* genes in bactobolin resistance (18). However, when sublethal bactobolin was added to the co-culture the competitive ability of the Δ*cdeAB-oprM* mutant increased by 1000-fold (Fig. 1). The increase was similar to that of wild-type *C. subtsugae*. These results show that although the CdeAB-OprM pump is important for bactobolin resistance, this pump is not responsible for the change in the outcome of competition following exposure to sublethal bactobolin.

We predicted a related pump with similar substrate specificity might be involved in the response to sublethal bactobolin in co-cultures. We searched the genome of CV017 for a gene related to *cdeB* and found one that is 80% identical to *cdeB* at the nucleotide level, which we call *cseB.* The *cseB* gene is flanked by genes with similarity to the *cdeA* and *oprM* genes (*cseA* and *oprN*) (Fig. 1B). Upstream of the *cdeAB-oprM* genes there is a gene encoding a putative TetR-family repressor called *cdeR*, which we previously showed is involved in regulating the *cdeAB-oprM* genes (18). However, there is no homologous gene to *cdeR* upstream of the *cseAB-oprN* gene cluster. Instead, downstream of the *cseAB-oprN* genes there is a gene encoding a predicted LysR family transcription activator, which we call *cseR* (21). To test the role of *cseAB-oprN* and *cseR* in our co-culture model, we deleted the *cseB* and *cseR* genes in CV017 and performed co-culture experiments with the deletion mutants. We observed that the competitive ability of both Δ*cseB* and Δ*cseR* was unchanged after exposure to sublethal bactobolin (Fig. 1, Δ*cseB* and Δ*cseR*, filled circles). Thus, both CseR and CseAB-OprN are important for *C. subtsugae* to increase competitiveness in response to sublethal bactobolin.

### Role of CseAB-OprN and CdeAB-OprM in antibiotic susceptibility and tolerance

To further characterize the newly identified CseAB-OprN and the CdeAB-OprM efflux pump, we determined the minimum inhibitory concentration (MIC) of bactobolin for *C. subtsugae* wild type and strains defective for each of the pumps or their putative regulators (Table 1). Consistent with previously reported results (18), the Δ*cdeAB-oprM* mutant was about 3-fold more susceptible to bactobolin than wild type. Deleting *cdeR* increased bactobolin resistance by 6-fold, and the increase was not observed in a *cdeR, cdeAB-oprM* double mutant. These results suggest CdeR is represses bactobolin resistance through the CdeAB-OprM system. We also observed *cdeA* and *cdeB* transcripts were increased in a *cdeR* mutant compared with wild type (Fig. S2). Together, these results support the idea that CdeAB-OprM increases bactobolin resistance and CdeR is a transcriptional repressor of the *cdeAB-oprM* gene cluster. In contrast, the bactobolin MIC for the Δ*cseB* mutant was identical to that of wild type (Table 1). Thus, the CseAB-OprN pump does not contribute to bactobolin resistance in standard MIC experiments.

**Table 1.** Antimicrobial susceptibility of *C. violaceum* (or C. *subtsugae*) strains.

| Cv strain | Minimum inhibitory concentration (MIC)<sup>a</sup> |  |  |  |  |  |  |  |
| --- | --- | --- | --- | --- | --- | --- | --- | --- |
|  | Bact<sup>b</sup><br>(%) | Tet | Gent | Kan | Ery | Chlo | Imip | Cipro |
| | | | | | $\mu\text{g.ml}^{-1}$ | | | $\text{ng ml}^{-1}$ |
| Wild type | 1.3±0.2 | 1.1±0.1 | 23±3 | 40 | 14±2 | 45.3±14 | 1.1±0.1 | 10±4 |
| $\Delta cdeAB^c$ | 0.4±0.2 | 0.5±0.1 | 23±3 | 38±4 | 7±1 | 12.5 | 1 | 2±1 |
| $\Delta cdeR$ | 8 | 5±1 | 36±5 | 41±4 | 75±10 | 106±31 | 1.1±0.1 | 110±12 |
| $\Delta cdeRAB^d$ | 0.5±0.1 | 0.4 | 45±5 | 47.5±3.5 | 45±5.8 | 45.3±14 | 1 | 350±70 |
| $\Delta cseB$ | 1.5±0.4 | 1.1±0.3 | 25±5 | 45±7 | 12±1 | 53±6 | 1 | 9±1 |
| $\Delta cseR$ | 1.4±0.4 | 1.0±0.3 | 25±5 | 35±7 | 16±1 | 54±7 | 1 | 8±1 |
| $\Delta cdeAB \Delta cseB$ | 0.3±0.1 | 0.4±0.2 | 26.7±3 | 37.5±4 | 6±2 | 54±7 | 1.1±0.1 | 2±1 |

<sup>a</sup>The minimum inhibitory concentration (MIC) of *B. thailandensis* culture fluid bactobolin, tetracycline (Tet), gentamicin (Gent), kanamycin (Kan), erythromycin (Ery), chloramphenicol (Chlo), and imipenem (Imip) and ciprofloxacin (Cipro). Results are the average of three independent experiments and the range is indicated when it was not zero.
<sup>b</sup>Results are from a single preparation of *B. thailandensis* fluid. Results with other preparations were similar. There were no observed growth defects in identical experiments with 3% culture fluid from a *B. thailandensis* bactobolin-deficient mutant.
<sup>c</sup> $\Delta cdeAB$ indicates the strain $\Delta cdeAB\text{-oprM}$
<sup>d</sup> $\Delta cdeRAB$ indicates the strain $\Delta cdeAB\text{-oprM}$ , $\Delta cdeR$

We also tested other antibiotics such as tetracycline, chloramphenicol, erythromycin (a macrolide), gentamicin and kanamycin (both aminoglycosides), imipenem (a cell wall-targeting β-lactam antibiotic), and ciprofloxacin (DNA gyrase inhibitor). In the case of tetracycline, erythromycin, chloramphenicol and ciprofloxacin, the *cdeAB-oprM* mutant was more susceptible than wild type. However, as with bactobolin, the Δ*cseB* mutant was not more susceptible than wild type to any of the tested antibiotics. We also tested a *cseB, cdeAB-oprM* double mutant for sensitivity to each of the antibiotics, and the double mutant showed the same susceptibility phenotypes as the *cdeAB-oprM* single mutant. The results show that CseAB-OprN does not play a significant role in increasing resistance in standard susceptibility experiments.

Although CseAB-OprN was not important in standard antibiotic susceptibility testing, the role of this pump in bactobolin-treated co-cultures (Fig. 1) supports the idea that this pump could be important for inducing a bactobolin tolerance response. To test this hypothesis, we developed an experiment to assess antibiotic tolerance by measuring the change in antibiotic susceptibility (MIC) following exposure to sublethal antibiotic concentrations. We grew CV017 cells for 6 h with 1/2 MIC of bactobolin (determined in Table 1), then compared the MIC of bactobolin-exposed cells with that of identically grown cells with no prior bactobolin exposure. We found that the MIC of bactobolin-exposed cells was 3.2 (± 0.2)-fold higher than identically treated cells not exposed to any antibiotic (Fig. 2A). Thus, we were able to observe induction of bactobolin tolerance in *C. subtsugae* in our experiment. We performed similar experiments with the *C. subtsugae* Δ*cdeAB-oprM*, Δ*cseB* or Δ*cseR* mutants. Our results showed exposure to sublethal bactobolin increased the MIC of the Δ*cdeAB-oprM* mutant 3.4 (± 1.2)-fold, similar to the induction observed with wild type. However, there was no significant MIC change observed with Δ*cseB* mutant cells following sublethal bactobolin exposure. These results support that the CseAB-OprN pump, and not the CdeAB-OprM pump, is important for bactobolin tolerance. Δ*cseR* mutant cells also induced bactobolin tolerance, but the change in MIC of 1.8 (± 0.2)-fold for the Δ*cseR* mutant was significantly less than that of wild type (*p* < 0.0005) supporting that CseR also plays a role in bactobolin tolerance.

**Fig. 2.**
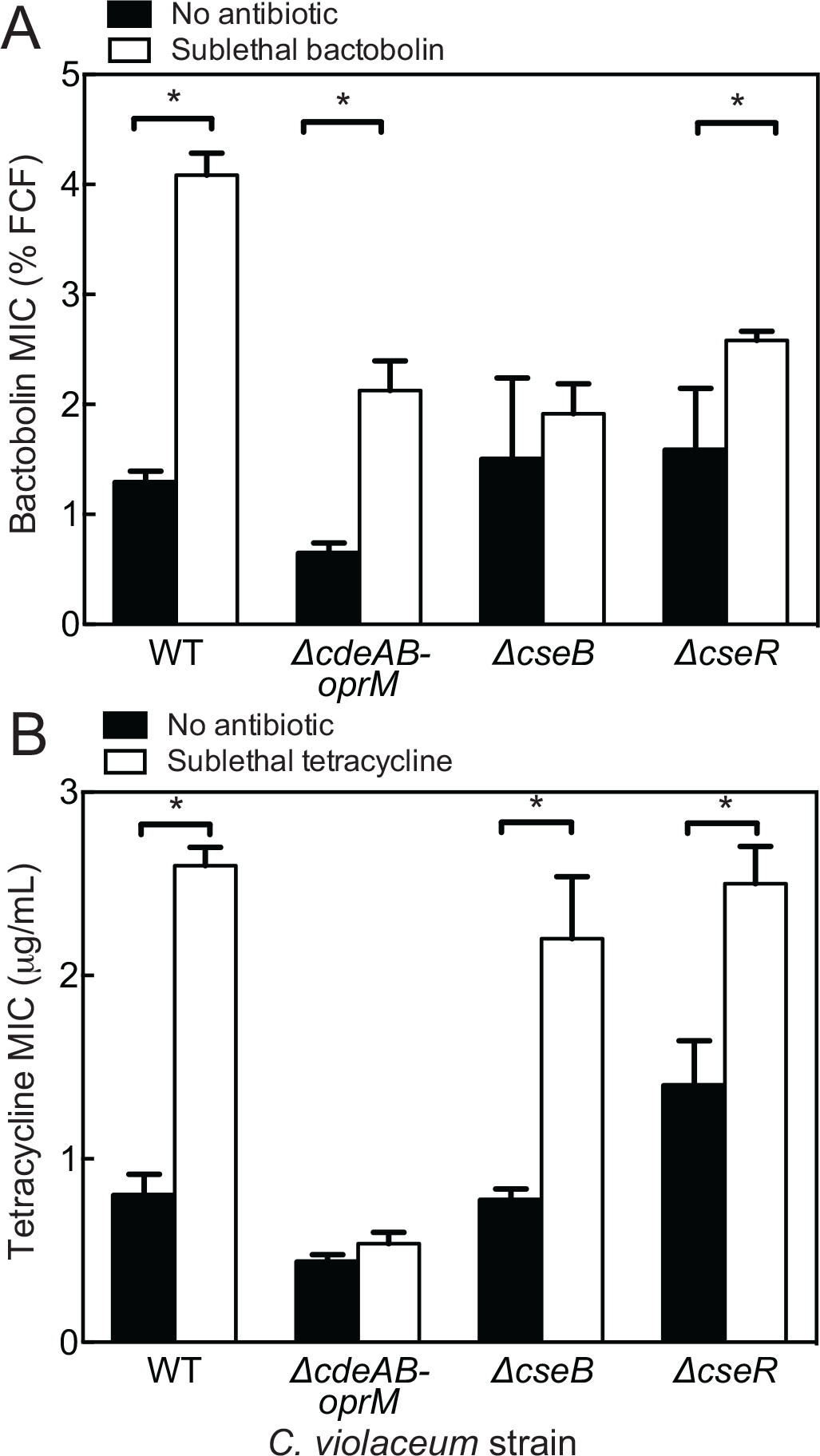
Antibiotic tolerance of *C. subtsugae* CV017 strains. Antibiotic tolerance is the change in minimum inhibitory concentration (MIC) of cells exposed to sublethal antibiotic (white bars) compared with that of identically treated cells with no antibiotic exposure (black bars). Sublethal antibiotic concentrations are 1/2 the MIC as indicated in Table 1 and used to treat cells for 6 h just prior to MIC determinations. Cells were treated for 6 h with A) Bactobolin or B) tetracyline and the same antibiotic was used for MICs. Bactobolin MIC is given as the final concentration of filtered culture fluid (% FCF). B) Tetracycline was used for sublethal exposure and MIC determinations. Final MIC is shown as the average and standard error of 4-6 biological replicates for each strain. Statistical significance by *t*-test; *, *P* < 0.03.

To test whether CseAB-OprN is involved in inducing resistance to antibiotics other than bactobolin, we used a similar approach to assess tolerance responses to erythromycin, ciprofloxacin, and tetracycline. We observed no significant change in MIC to erythromycin or ciprofloxacin after sublethal exposure (Fig. S3). However, exposure to sublethal concentrations of tetracycline increased tetracycline MIC by 3.9 (± 0.9)-fold, similar to that of bactobolin (Fig. 2B). Surprisingly, a similar 2.7 (± 1.3)-fold increase in MIC was observed for the Δ*cseB* mutant. And no increase was observed for the Δ*cdeAB-oprM* mutant. These results show *C. subtsugae* induces tetracycline tolerance through the CdeAB-OprM system and not through CseAB-OprN.

### Specificity of CseAB-OprN and CdeAB-OprM

We found it interesting that the CdeAB-OprM system was important for bactobolin resistance, but not tolerance (Fig. 2). These results suggest bactobolin might be weakly recognized as an inducer by the CdeAB-OprM system or serve as a poor substrate for export. To understand the interaction of each pump with bactobolin, we exposed cells to sublethal concentrations of bactobolin and then determined the MIC of tetracycline. We predicted that if bactobolin can induce the CdeAB-OprN pump but serves as a poor substrate for export, we would observe an increase in tetracycline MIC in response to bactobolin induction in a manner dependent on CdeAB-OprM. Consistent with this prediction, our result showed that incubation with sublethal bactobolin increased the tetracycline MIC and the response was dependent on CdeAB-OprM (Fig. 3A). These results support the conclusion that the CdeAB-OprM pump can be induced by sublethal bactobolin. We also performed the reverse experiment; we incubated cells with sublethal tetracycline and subjected those cells to a bactobolin MIC. In this experiment, tetracycline increased the bactobolin MIC in a manner that was also dependent on the CdeAB-OprM system (Fig. 3B). Thus, tetracycline and bactobolin can both serve to induce tolerance through CdeAB-OprM, although the effects of bactobolin were only observed in combination with tetracycline suggesting bactobolin is a relatively poor substrate of this pump. In contrast, tetracycline was unable to induce tolerance or be recognized by the CseAB-OprN pump, suggesting this antibiotic is only recognized by the CdeAB-OprM pump. Together, our results support the conclusion that CseAB-OprN has a high level of sensitivity and specificity for bactobolin, whereas CdeAB-OprM recognizes a broader range of antibiotics.

**Fig. 3.**
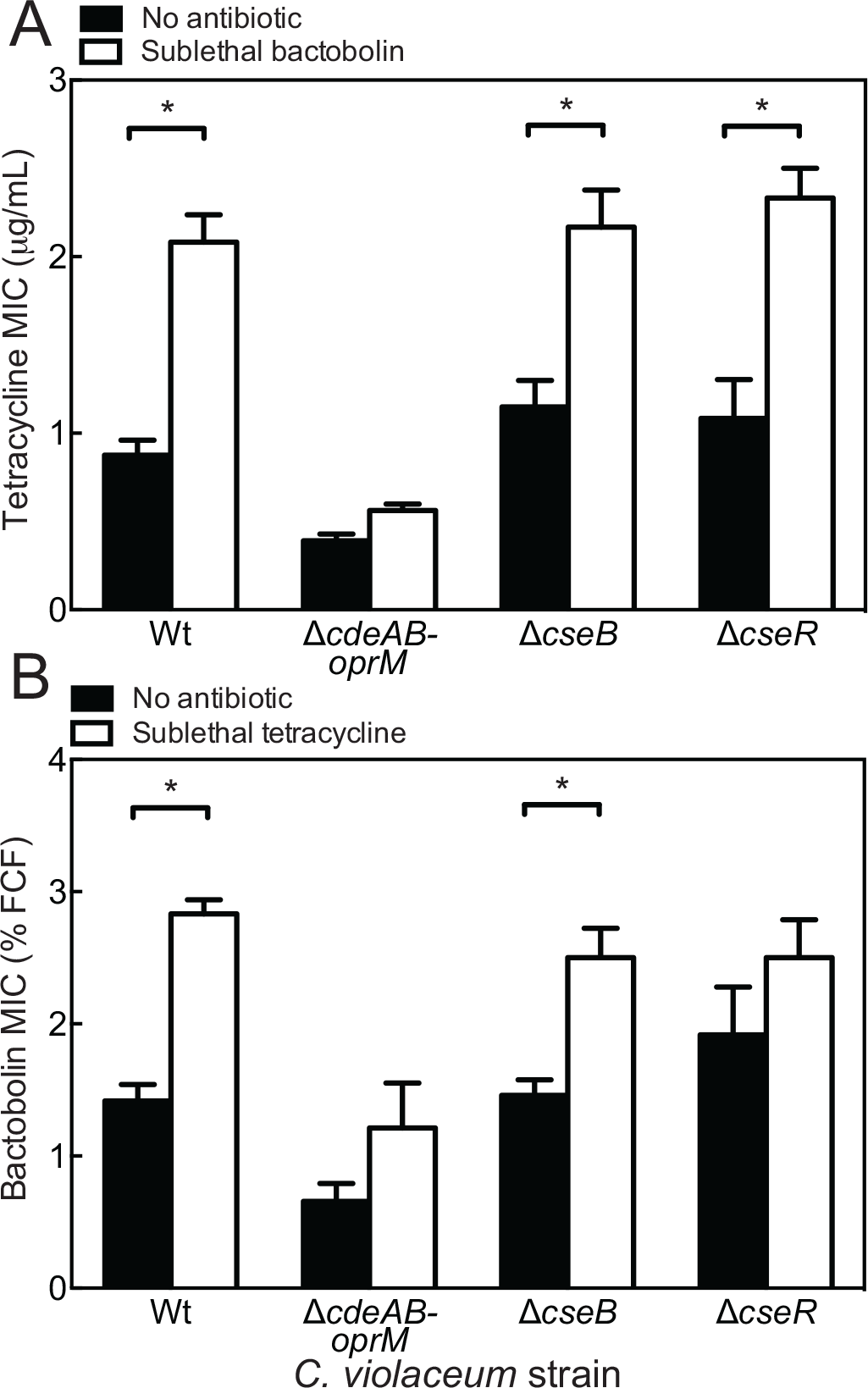
Specificity of tolerance responses. Antibiotic tolerance experiments were performed as described for Fig. 2. In this case, sublethal exposure and MIC determinations were with different antibiotics: A) bactobolin and tetraycline or B) tetraycline and bactobolin, respectively. Final MIC is shown as the average and standard error of 3-4 biological replicates for each strain. *, statistical significance by *t*-test; *, *P* < 0.03.

### CseR is a bactobolin-responsive *cseAB-oprN* gene activator

Because CseR is important in co-culture and antibiotic tolerance experiments and also belongs to the LysR transcriptional regulator family, we hypothesized that CseR might be an activator of the CseAB-OprN system. To test this hypothesis, we measured *cseA* transcripts in wild type or the Δ*cseR* mutant cells exposed to bactobolin, tetraycline or no antibiotic (Fig. 4). Our results showed a small but significant 2.4-fold increase in *cseA* transcripts in bactobolin-treated cells compared with untreated cells or tetracycline-treated cells. The increase in bactobolin-treated cells was dependent on *cseR.* Thus, CseR activates transcription of the *cseAB-oprN* genes in response to bactobolin. We also measured induction of *cdeA* transcription in these cells to assess the specificity of CseR gene activation. We observed a 3-fold increase in *cdeA* transcription in cells treated with either bactobolin or tetracycline that was independent of CseR. Thus, CseR specifically activates *cseAB-oprN* transcription in response to bactobolin antibiotic. We also observed that *cdeA* transcription was induced by bactobolin or tetracycline in a CseR-independent manner, which was consistent with the results of our antibiotic tolerance experiments (Figs. 2 and 3).

**Fig. 4.**
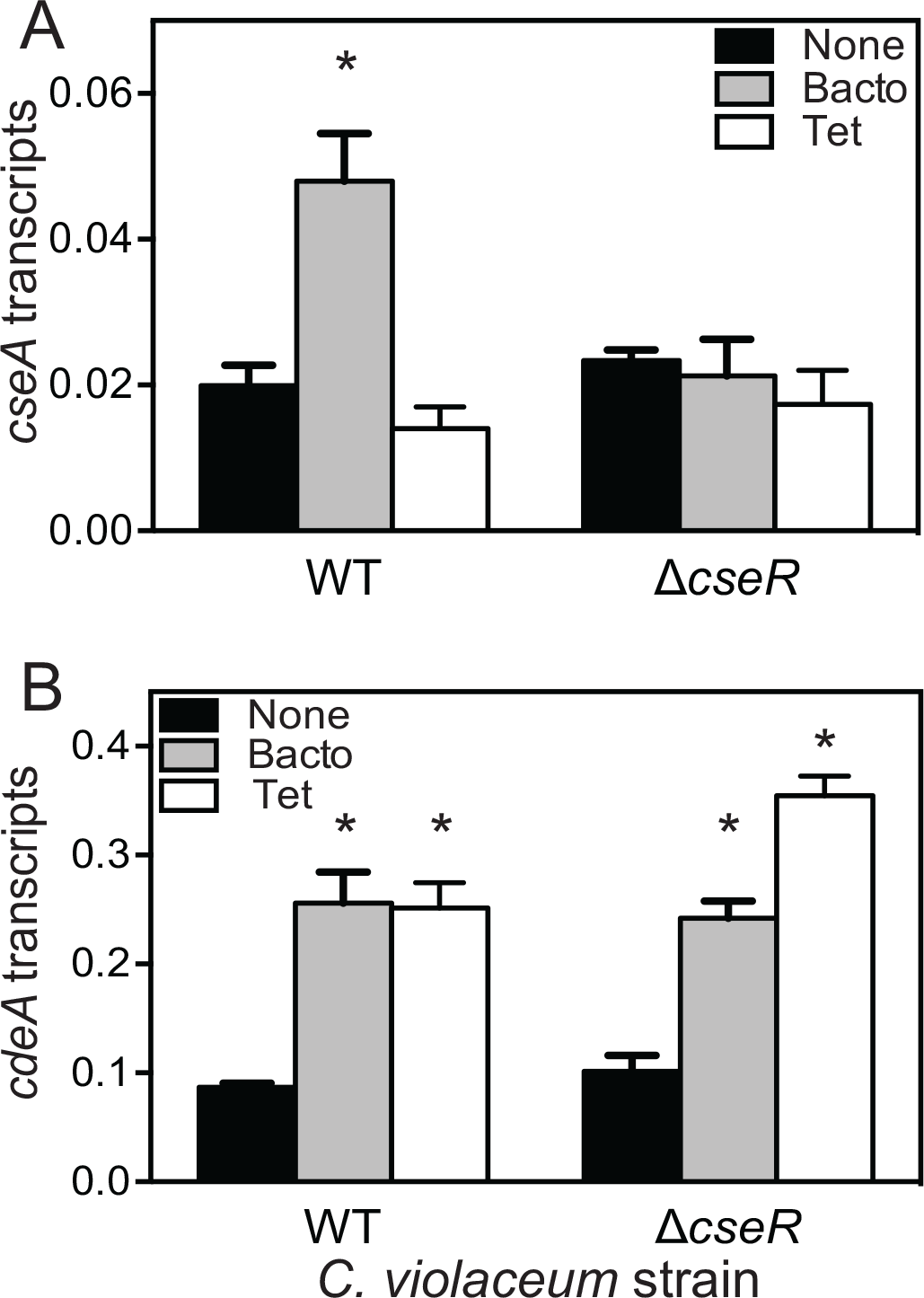
Antibiotic induction of the efflux pump genes. *cseA* (A) or *cdeA* (B). Transcripts were measured from exponentially growing cells treated for 4 h with no antibiotic (black bars) or a sublethal concentration of bactobolin (grey bars) or tetracycline (white bars). Sublethal antibiotic concentrations are 1/2 the MIC as indicated in Table 1. Results are shown as *rpoD-*adjusted transcripts. The values represent the mean of three independent experiments and the vertical bars represent the standard error of the mean. Statistical analysis by t- test compared with untreated wild type: *, p ≤ 0.02

Many LysR-family activators require a ligand for activation of target genes. To test if increasing CseR production is sufficient to activate the CseAB-OprN system or if bactobolin is required for this activation, we introduced the *cseR* gene into plasmid pBBR-MCS5 downstream of the constitutive T3 promoter and introduced this plasmid to the *C. subtsugae* Δ*cseR* strain. As a control, we also made a similar pBBR-MCS5 *cdeR* plasmid and introduced this plasmid to the *C. subtsugae* Δ*cdeR* strain. While the pBBR-*cdeR* plasmid markedly decreased the Δ*cdeR* mutant bactobolin MIC, there were no effects of expressing *cseR* in the Δ*cseR* mutant (Table S1). However, *cseR*-expressing cells exposed to sublethal bactobolin significantly increased bactobolin tolerance compared with the plasmid-only control or cells not exposed to bactobolin (Fig. S3). Thus, constitutive CseR production does not appear to be sufficient to induce tolerance, and bactobolin likely serves as a ligand activator of the CseR regulator.

### Distribution of *cdeAB-oprM* and *cseAB-oprN* in *Chromobacterium*

We were interested in evaluating the distribution of the *cdeAB-oprM* and *cdeAB-oprM* genes in the *Chromobacterium* genus. We constructed a phylogenetic tree using 32 *Chromobacterium* genomes and an outgroup (*Aquitaliea magnus*) (Fig. 5, Fig. S5 and Table S2). We constructed the tree using a set of 140 orthologous proteins (see Materials and Methods) to increase confidence of relationship predictions. Using this method, CV017 grouped with the *C. subtsugae* strains, consistent with the new classification of this strain. We were also able to make confident predictions of species associations for some of the unnamed species, for example, strain F49 grouped with the *C. subtsugae* clade. It is notable that the species known to cause infections in humans, *C. violaceum* and *C. haemolyticum*, grouped separately from *C. subtsugae*. Thus *C. subtsugae* may be representative of a primarily non-pathogenic clade of *Chromobacterium.*

**Fig. 5.**
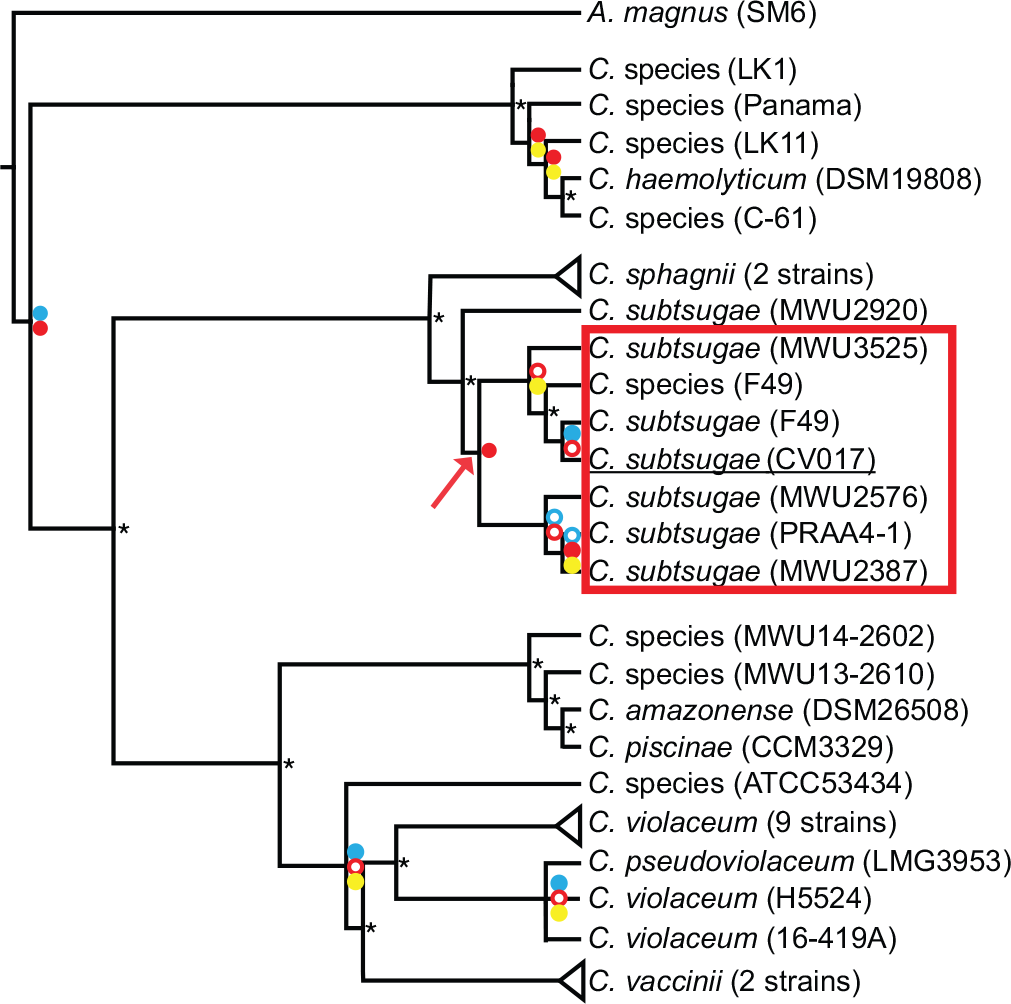
Phylogenetic tree of *Chromobacterium* species. Tree construction is described in materials and methods. Circles at each node represent bootstrap values based on neighbor joining (blue circles), maximum likelihood (red circles) and Bayesian (yellow circles) methods. Filled circles, >95% support; unfilled circles, >50% support; missing circles, <50% support. *, all three methods showed >95% support. Triangles represent several highly similar strains; *Chromobacterium sphagnii* (37-2 and 14-B11), *Chromobacterium violaceum* (L_1B5_1, LK17, LK15, GHS51, 16-454, CV1192, CV1197, GN5 and 12472), and *C. vaccinii* (MWU205 and 21-1). A full representation of the species in the phylogenetic tree is shown in Fig. S1. Red box indicates species encoding the *cseAB-oprN* genes. Red arrow, node where species with the *cseAB-oprN* gene cluster separate from other species.

We identified *cdeB* homologs in all of the strains ranging from 85-100% nucleotide sequence identity to CV017 *cdeB*. Each of these strains had a similar arrangement of *cdeA, oprM* and *cdeR* genes surrounding *cdeB* (Fig. 5B). We used a best-reciprocal BLAST search to discern *cdeB* and *cseB* homologs, and identified *cdeB* homologs only in CV017, F49 and other members of the *C. subtsugae* species with the exception of MWU2920. In these other species, the *cseB* genes were >99% identical to one another, and the organization of genes surrounding *cseB* was conserved with that of the *cseAB-oprN* genes in CV017. We predicted that the *cseAB-oprN* gene cluster might have arisen from the *cdeAB-oprM* genes via a duplication event that occurred in an ancestor of *C. subtsugae*; however, generation of a second tree using the *cseB* and *cdeB* gene homologs did not support this hypothesis (Fig. S6). Instead, the alignments show *cseB* groups with *cdeB* from *Chromobacterium* species C61, suggesting that *cseB* might have arisen from a horizontal gene transfer event from an ancestor of this strain.

## DISCUSSION

In this study, we demonstrate antibiotic specificity and regulation of two efflux pumps of *Chromobacterium subtsugae* CV017. One of these, CseAB-OprN, had not been previously reported and is limited to the *C. subtsugae* species. The other, CdeAB-OprM, is widely distributed in members of the *Chromobacterium* genus. CdeAB-OprM has a broad range of specificity for antibiotics such as tetracycline, erythromycin, chloramphenicol and ciprofloxacin. As ciprofloxacin is one of the recommended treatments for infections with pathogenic *Chromobacterium* (2), the results suggest CdeAB-OprM might be important in cases of antibiotic failures during *Chromobacterium* infections. To date, there are very few publications on *Chromobacterium* mechanisms of antibiotic resistance and drug export (18, 22, 23). Results of this study will be useful for identifying resistance mechanisms in clinical isolates and for future *Chromobacterium* research. As *Chromobacterium* members are primarily saprophytic non-pathogens, our results are also important for understanding the ecology and evolution of efflux pumps.

Our results demonstrate that both CdeAB-OprM and CseAB-OprN are induced by their antibiotic substrates. Such a response is typical of related pumps such as the aminoglycoside-responsive MexXY pump of *Pseudomonas aeruginosa* (24) and the trimethoprim-responsive BpeEF-OprC pump of *Burkholderia pseudomallei* (25). Because many antibiotic resistance factors are costly to produce, waiting to induce their production until needed might allow a valuable metabolic cost savings (26). The ability to respond to antibiotic stress might be an important strategy for bacteria to survive antibiotics during treatment of infected patients (27). Such responses might also be important during competition with other bacteria (28). Our co-culture model highlights the importance of tolerance induction during competition and how such regulation might evolve in soil bacteria. The induction of tolerance has been proposed to be a type of ‘danger sensing,’ which is important for sensing and responding to threats posed by other bacteria (28). Sublethal antibiotics serves as a warning that antibiotic-producing competitors are nearby and might soon deliver higher killing doses.

Competitive interactions with other bacteria might also be an important factor shaping the evolution of antibiotic production. Many antibiotics are regulated by quorum sensing, which activates production in response to changes in population density; for example in *B. thailandensis*, bactobolin is regulated by the BtaR2-I2 quorum-sensing system (29). Placing antibiotics under the control of quorum-sensing systems might be a strategy to avert induction of antibiotic tolerance by other species (30). Quorum-sensing control of antibiotics might serve to delay antibiotic production until the population can produce a sudden killing dose of antibiotic. Bactobolin production is activated by quorum sensing when the population reaches stationary phase (31) and sufficient cells have accumulated to produce killing dose. Adding bactobolin at the start of the experiment exposes *C. substugae* to low doses prior to the accumulation of the killing dose. The early exposure allowed *C. subtsugae* to mount a defense response and survive the higher killing doses. Thus regulatory mechanisms that delay antibiotic production might increase the killing effect of the antibiotics, allowing an effective and efficient ‘sneak attack’ on the competitor (30).

Our phylogenetic analysis suggests that the *cseAB-oprN* genes were acquired through horizontal transfer of the *cdeAB-oprM* cluster from another species, rather than through duplication of the *cdeAB-oprM* genes in the ancestral strain (Fig. 5A and S6). The finding that horizontal transfer of resistance genes occurred in saprophytic species supports the idea that horizontal transfer can lead to the dissemination of antibiotic resistance genes in polymicrobial communities of the soil (32). Our co-culture experiments also support the idea that the spread of resistance might have been helpful for surviving antibiotics produced by other bacteria in the soil. The acquisition of new resistance genes might be important for surviving antibiotics produced by other species, as demonstrated by our co-culture model, or it might also promote tolerance to self-produced antibiotics (33). *Chromobacterium* is known to produce an arsenal of antimicrobial-active molecules (34), although many have not been well characterized. The hypothesis that CseAB-OprN or CdeAB-OprM provides resistance to a self-produced antibiotic can be tested experimentally.

## MATERIALS AND METHODS

### Bacterial culture conditions and strains

All strains were grown in Luria-Bertani (LB) broth, LB containing morpholine-propanesulfonic acid (LB-MOPS; 50 mM; pH 7), or on LB with 1.5% (wt/vol) Bacto-Agar. All broth cultures were incubated with shaking at 30°C (*C. subtsugae* or *C. subtsugae-B. thailandensis* co-cultures) or 37°C (*B. thailandensis* or *E. coli*). As a source of bactobolin, we used filtered *B. thailandensis* culture fluid prepared as described previously (18). Filtered *B. thailandensis* culture fluid was stored at 4°C for up to 1 month prior to use and used directly for experiments. For strain constructions, we used gentamicin at 50 μg ml−1 (*C. subtsugae*) or 15 μg ml−1 (*E. coli*). For selection from co-cultures we used gentamicin at 100 μg ml−1 (*B. thailandensis*) and trimethoprim at 100 μg ml−1 (*C. subtsugae*).

Bacterial strains and plasmids used in this study are listed in Tables S3-S4. For co-cultures, we used the wild-type *B. thailandensis* strain E264 (35). *C. subtsugae* strain CV017 (referred to as wild type, previously known as *C. violaceum* CV017) is a derivative of strain ATCC31532 (36) with a transposon insertion in gene CV_RS05185 causing overexpression of violacein (37, 38). All *C. subtsugae* mutant strains were constructed from CV017 using allelic exchange and methods described previously (16). The constructs for generating deletions in *cseB, cdeR* and *cseR* were made by introducing PCR-generated amplicons or synthetic gene fragments (IDT or Genscript) into pEX18Gm-derived delivery plasmid (39). The mutation was introduced to *C. subtsugae* by conjugation and transconjugants were selected on gentamicin. Transconjugants were transferred to no salt-LB + 15 % sucrose (wt/vol) to select for plasmid excision and correct clones were identified by PCR-based screening. *cseR* and *cdeR* expression plasmids were made using pBBR-MCS5 (40) with PCR-generated *cseR* or *cdeR* fragments introduced downstream of the T3 promoter. All strains and PCR-generated plasmids were verified by PCR amplification and sequencing.

### Co-culture experiments

Co-culture experiments were conducted in 10 ml LB-MOPS medium in 125 ml baffled flasks. The inoculum was from logarithmic-phase pure cultures of *C. subtsugae* and *B. thailandensis*. The initial OD_600_ in the co-culture was 0.05 for *B. thailandensis* (2-4×10^7^ cells per ml) and 0.005 for *C. subtsugae* (2-4×10^6^ cells per ml). When indicated, cultures also contained *B. thailandensis* culture fluid at a concentration corresponding with ½ of the MIC for the *C. subtsugae* strain used in the experiment. After inoculating, co-cultures were incubated at 30°C with shaking at 230 r.p.m for 24 h. Colony forming units of each species were determined by using differential antibiotic selection on LB agar plates. *B. thailandensis* was selected with gentamicin and *C. subtsugae* was selected with trimethoprim.

### Sequencing the *cse* gene locus

*cseB* (CV_15400) was initially identified in the assembled CV017 genome (GCA_001510755.1) on scaffold 169 (LKIW01000078). 274 bases downstream of *cseB* is the 3’ end of scaffold 169. As this sequence did not overlap with any of the other 211 scaffolds of CV017, we used a BLAST search to identify similar sequence in a similar strain (*C. subtsugae* PRAA4-1, GCA_001676875.1) This search identified sequence containing a *cseB-*like gene in scaffold 30 of PRAA4-1, corresponding with CV017 scaffold 192 (LKIW01000104). We used alignments of CV017 scaffold 169 and 192 with the PRAA4-1 sequence to design oligonucleotides to amplify the region between scaffolds 169 and 192. We Sanger sequenced the amplified PCR product to verify the correct orientation of the scaffolds and identify the previously unknown 39 nucleotides linking forward-facing scaffold 169 with reverse-facing scaffold 192. This was GCCGCCGCCGACAGCCAGCGCCAGCGCGTCAGCGCGACG. CV017 scaffold 169 was updated through NCBI (LKIW01000078.2) and scaffold 192 was made secondary to this new scaffold so searches for either will give the new version of scaffold 169.

### Antibiotic susceptibility and tolerance experiments

Antibiotic susceptibility (Table 1) was determined by minimum inhibitory concentration (MIC) according to the 2003 guidelines of the Clinical and Laboratory Standards Institute (NCCLS), using a modified microtiter method. Antibiotics were added to LB-MOPS in a 100 ul well of a 96-well plate, and 10 successive 2-fold dilutions were made. For each antibiotic, 3 different starting concentrations of antibiotic were used to increase sensitivity of the experiment. *C. subtsugae* inocula were prepared by diluting logarithmic-phase cells from LB-MOPS cultures to an optical density at 600 nm (OD_600_) of 0.005 (6×10^6^ CFU) in each well. Plates were sealed with Breathe-Easy strips (Fisher Scientific, fishersci.com) and grown for 24 h with shaking at 30°C. The MIC was defined as the lowest concentration of antibiotic (μg/ml) in which bacterial growth in the well was not measurable by determining the optical density at 600 nm (OD_600_) on a 96-well plate reader.

To assess changes in MIC induced by exposure to sublethal antibiotic (tolerance), logarithmic-phase *C. subtsugae* cells were diluted to an OD_600_ of 0.1 in 10 ml LB-MOPS in a 125-ml culture flask. The culture medium contained antibiotic at ½ of the MIC determined by the method described above (this varied for each strain), or no antibiotic (untreated), and incubated for 6 hours with shaking at 30°C. These cultures were then directly used to determine MIC using methods described above.

### Droplet digital PCR

RNA was harvested from *C. subtsugae* as described in the figure legends. RNA was prepared using the RNeasy^®^ Mini Kit (Qiagen) following the manufacturer instruction with a modification in the DNA digestion step as described previously (18). Droplet digital PCR was performed on a Bio-Rad’s QX200 Droplet Digital PCR (ddPCR) System using Eva Green Supermix. Each reaction mixture contained 1 ng μl^−^1 of cDNA template, 0.25 μM of each primer, 10 μl Eva Green Supermix, and 8 μl H_2_O in a 20-μl volume. After generating 40 μl of oil droplets, 40 rounds of PCR were conducted using the following cycling conditions: 94°C for 20 sec, 60°C for 20 sec, and 72°C for 20 sec. Absolute transcript levels were determined using the Bio-Rad QuantaSoft Software. In all cases a no-template control was run with no detectable transcripts.

### Phylogeny of *Chromobacterium* species and efflux pump genes

*Chromobacterium* species used for phylogenetic analyses are listed in Table S4. Annotated protein sequences from assembled genomes of *Chromobacterium* and *Aquitaliea magnusonii* (outgroup) were retrieved from the National Center for Biotechnology Information (NCBI, June 2018, Table S3). Comparisons using 16S rRNA provided low confidence predictions of phylogeny, thus we performed an analysis using a set of 140 single-copy orthologs, which were identified using several steps. Orthologous proteins were initially identified by using blastp to carry out reciprocal best BLAST hits of each protein from each strain against a protein database made of all the proteins in our strain set ((41), options: – max_target_seqs 1), to find orthologs with best BLAST hit between all possible pairs of species, and no more than 10% variation in protein length. We aligned this group of 171 proteins individually using Muscle version 3.8.31 ((42), options -diags), then reordered sequences using the stable.py script provided by the Muscle developer. Finally, we removed any orthologous protein groups with less than 75% average pairwise identity, less than 30% of sites identical or 100% of sites identical. This last step ensured that all proteins would have intermediate levels of divergence and left the final set of 140 orthologs used for phylogenetic tree construction. Finally, we concatenated the protein sequences in each alignment to create one single alignment consisting of 25,351 amino acids. After inspection of this alignment, we found that several pairs of isolates had very low divergence (less than 10 amino acid differences across the entire alignment). We decided to remove one of each of these pairs to reduce redundancy (removed isolates were *C. violaceum* strains LK6, LK30, H5525 and 16-419B). C. *subtsugae* F49 and *C. subtsugae* CV017 are identical across the alignment of orthologous proteins but both were retained because CV017 is the focus of the paper.

We generated phylogenetic trees using neighbor joining, maximum likelihood and Bayesian methods. A simple neighbor joining (NJ) tree implemented in Geneious version 10.1.3 (http://www.geneious.com) with a Jukes-Cantor substitution model and 100 bootstrap replicates and *A. magnusonii* as an outgroup. We constructed a maximum likelihood tree using RaxML version 8.2.11 (43) with a Gamma BLOSSUM62 protein model, 100 bootstrap replicates with a parsimony random seed of 1 and *A. manusonii* as an outgroup. We used Mr. Bayes version 3.2.6 (44) to construct a tree using Bayesian methods with a Poisson rate matrix, gamma rate variation with 4 categories, and *A. manusonii* as an outgroup, an initial chain length of 1,100,000 with four heated chains at a temperature of 0.2, the subsampling frequency was 2000 generations after an initial burn-in length of 100,000 generations. However, after 332,000 generations, the standard deviation of split frequencies was less than 0.01 and so the tree search was terminated. For all three methods, consensus trees were built based on 50% majority rule, and the three trees were compared using the RaxML maximum likelihood tree as a backbone. We elected to use a concatenated sequence to build our tree because our concern is about the overall evolutionary history of the genus and not how specific genes trees might differ from species tree.

To reconstruct the evolutionary history of the two efflux pumps, we compared DNA sequence of the pump genes to improve our ability to resolve evolutionary distances between closely related isolates. We retrieved the DNA sequences by performing a tblastn (45) of the amino acid sequence for that protein against all *Chromobacterium* genome assemblies and then located the correct sequence based on isolate name. Note that one sequence was split among two scaffolds in *C. subtsugae* MWU2387 and was therefore not annotated so we pieced together the appropriate sequence. To find orthologs of the second pump (*cseB*) we performed a tblastn search of the amino acid sequence of CdeB from CV017 against all assembled *Chromobacterium* species. Reciprocal best BLAST searches confirmed the presence of this second pump in only a subset of the *C. subtsugae* isolates (including CV017). We also searched for similar sequences outside of *Chromobacterium* to find evidence for horizontal transfer of the pump. Finally, we constructed phylogenetic trees of the DNA sequence for the two pumps similar to what was described above but with nucleotides instead of amino acids.

## ACKNOWLEDGEMENTS

This work was supported by a startup package and a Graduate Research Fund from the University of Kansas and by an NIH COBRE Center for Molecular Analysis of Disease Pathways Research Project Award to J.R.C. (P20GM103638). S.B. was supported by an NIH K-INBRE fellowship (P20GM103418) and K.C.E. was supported by the NIH Chemical Biology Training Program (T32 GM08545). R.L.U. was supported by the NIH (grants R00-GM114714 and R01-AI139154-01). We thank the KU K-INBRE Bioinformatics Core (P20GM103418) and Stuart Macdonald for helpful input on sequence analysis.

