## Supplemental File for "Efflux pumps in *Chromobacterium* species and their involvement in antibiotic tolerance and survival in a co-culture competition model"

**Table S1. Antimicrobial susceptibility of *C. subtsugae* strains expressing *cdeR* and *cseR.***

| | Minimum inhibitory concentration (MIC)[a] | |
|---|---|---|
| | Bact[b] | Tet |
| *Cv* strain | (%) | $\mu g.ml^{-1}$ |
| Cv017 pBBRMCS-5 | 1.88±0.6 | 1.69±0.7 |
| Cv017Δ*cseR* pBBRMCS-5 | 1.38±0.4 | 1.56±0.5 |
| Cv017Δ*cseR* pBBR*cseR* | 1.63±0.4 | 1.63±0.4 |
| Cv017Δ*cdeR* pBBRMCS-5 | 8.00±0.0 | 5.75±1.5 |
| Cv017Δ*cdeR* pBBR*cdeR* | 1.31±0.5 | 0.88±0.3 |

[a]The minimum inhibitory concentration (MIC) of *B. thailandensis* culture fluid bactobolin, and tetracycline (Tet). Results are the average of four independent experiments and the range is indicated when it was not zero.

[b]Results are from a single preparation of *B. thailandensis* fluid.

**Table S2. Bacterial strains used for phylogenetic analysis (Fig. 5, S4 and S5).**

| Species | Strain | Assembly accession |
|---|---|---|
| *Aquitaliea magnusonii* | SM6 | |
| *Chromobacterium* spp. | ATCC53434 | |
| *Chromobacterium* spp. | C-61 | |
| *Chromobacterium* spp. | F49 | |
| *Chromobacterium* spp. | MWU13-2610 | |
| *Chromobacterium* spp. | MWU14-2602 | |
| *Chromobacterium* spp. | LK1 | |
| *Chromobacterium* spp. | LK11 | |
| *Chromobacterium* spp. | Panama | |
| *Chromobacterium amazonense* | DSM26508 | |
| *Chromobacterium haemolyticum* | DSM19808 | GCA_000711885.1 |
| *Chromobacterium piscinae* | CCM3329 | |
| *Chromobacterium pseudoviolaceum* | LMG395 | |
| *Chromobacterium sphagnii* | 14-B11 | GCA_001855555.1 |
| *Chromobacterium sphagnii* | 37-2 | GCA_001855575.1 |
| *Chromobacterium subtsugae* | F49 | GCA_000812805.1 |
| *Chromobacterium subtsugae* | MWU2387 | GCA_001020525.1 |
| *Chromobacterium subtsugae* | MWU2576 | GCA_001020515.1 |
| *Chromobacterium subtsugae* | MWU2920 | GCA_001020585.1 |
| *Chromobacterium subtsugae* | MWU3525 | GCA_001020505.1 |
| *Chromobacterium subtsugae* | PRAA4-1 | GCA_001676875.1 |
| *Chromobacterium subtsugae* | CV017 | GCA_001510755.1 |
| *Chromobacterium vaccinii* | 21-1 | GCA_001855275.1 |
| *Chromobacterium vaccinii* | MWU205 | GCA_000971335.1 |
| *Chromobacterium violaceum* | 16-419A | GCA_002081735.1 |
| *Chromobacterium violaceum* | 16-454 | GCA_002081775.1 |
| *Chromobacterium violaceum* | ATCC12472 | GCA_000007705.1 |
| *Chromobacterium violaceum* | CV1192 | GCA_002735945.1 |
| *Chromobacterium violaceum* | CV1197 | GCA_002735965.1 |
| *Chromobacterium violaceum* | GHPS1 | GCA_002179535.1 |
| *Chromobacterium violaceum* | GN5 | GCA_000812485.1 |
| *Chromobacterium violaceum* | H5524 | GCA_002081875.1 |
| *Chromobacterium violaceum* | L_1B5_1 | GCA_000952105.1 |
| *Chromobacterium violaceum* | LK15 | GCA_001043755.1 |
| *Chromobacterium violaceum* | LK17 | GCA_001043735.1 |

**Table S3. Bacterial strains used in this study.**

| Strain | Relevant properties | Reference or source |
|---|---|---|
| *Chromobacterium subtsugae* | | |
| CV017 | Mini-Tn5 mutant of ATCC 31532, ‘wild type’ | (1) |
| CV017 Δ*cdeAB* | CV017 containing an unmarked, in-frame *cdeAB-oprM* deletion | (2) |
| CV017 Δ*cseB* | CV017 containing an unmarked, in-frame *cseB* deletion | This study |
| CV017 Δ*cdeR* | CV017 containing an unmarked, in-frame *cdeR* deletion | This study |
| CV017 Δ*cseR* | CV017 containing an unmarked, in-frame *cseR* deletion | This study |
| CV017 Δ*cdeRAB* | CV017 Δ*cdeAB-oprM* containing an unmarked, in-frame *cdeR* mutation | This study |
| CV017 Δ*cdeAB* Δ*cseB* | CV017 *cdeAB-oprM* containing an unmarked, In-frame *cseB* mutation | This study |
| *Burkholderia thailandensis* | | |
| E264 | Wild-type strain | (3) |
| BD20 | *btaK* (bactobolin) mutant of E264 | (4) |
| *Escherichia coli* | | |
| DH5α | [F- φ80 lacZΔM15] Δ(lacZYA-argF)U169 recA1 endA1 hsdR17, [rK−mK+] supE44 thi-1 gyrA relA1 | Invitrogen |
| Rho-3 | thi-1 thr-1 leuB26 tonA21 lacY1 supE44 recA integrated RP4-2 Tcr::Mu (λpir+)Δasd::FRT ΔaphA::FRT | (5) |

**Table S4. Bacterial plasmids used in this study.**

| Plasmid | Relevant properties | Reference or source |
|---|---|---|
| pEX18Gm | Suicide vector, $Gm^r$ | (6) |
| pEX18Gm Δ*cdeAB-oprM* | pEX18Gm with Δ*cdeAB-oprM* extending from +22 of *cdeA* to +1362 of *oprM* with regard to translation start site | (2) |
| pEX18Gm Δ*cdeR* | pEX18Gm with Δ*cdeR* extending from +21 to +624 with regard to translation start site | This study |
| pEX18Gm Δ*cseB* | pEX18Gm with Δ*cseB* extending from +43 to +3141 with regard to translation start site | This study |
| pEX18Gm Δ*cseR* | pEX18Gm with Δ*cseR* extending from +13 to +885 with regard to translation start site | This study |
| pBBRmcs5 | Broad-host-range vector; $Gm^r$ | (7) |
| pBBR *cseR* | pBBRmcs5 with *cseR* extending from +498 to +873 with regard to translation start site | This study |
| pBBR *cdeR* | pBBRmcs5 with *cdeR* extending from +458 to +1114 with regard to translation start site | This study |

**Fig. S1. *B. thailandensis-C. subtsugae* competition.** Co-cultures were of wild-type *B. thailandensis* (*Bt*) and *C. subtsugae* strain CV017 (*Cs*). The dashed line indicates the starting 1:10 ratio of *Cs* to *Bt.* The ratio of *Cs* to *Bt* was determined after 24 h by selective plating and colony counts. Open circles, co-cultures with no additions. Filled circles, co-cultures with sublethal bactobolin provided by adding *Bt* culture fluid (0.6%) at the start of the co-culture experiment, or with 1% fluid from a *Bt* bactobolin mutant (BD20). The solid lines represent means for each group. The vertical bars show the standard error of the mean for each group. *****, statistically significant by student's *t*-test compared with untreated or treated with bactobolin-deficient culture fluid ($p < 0.05$).

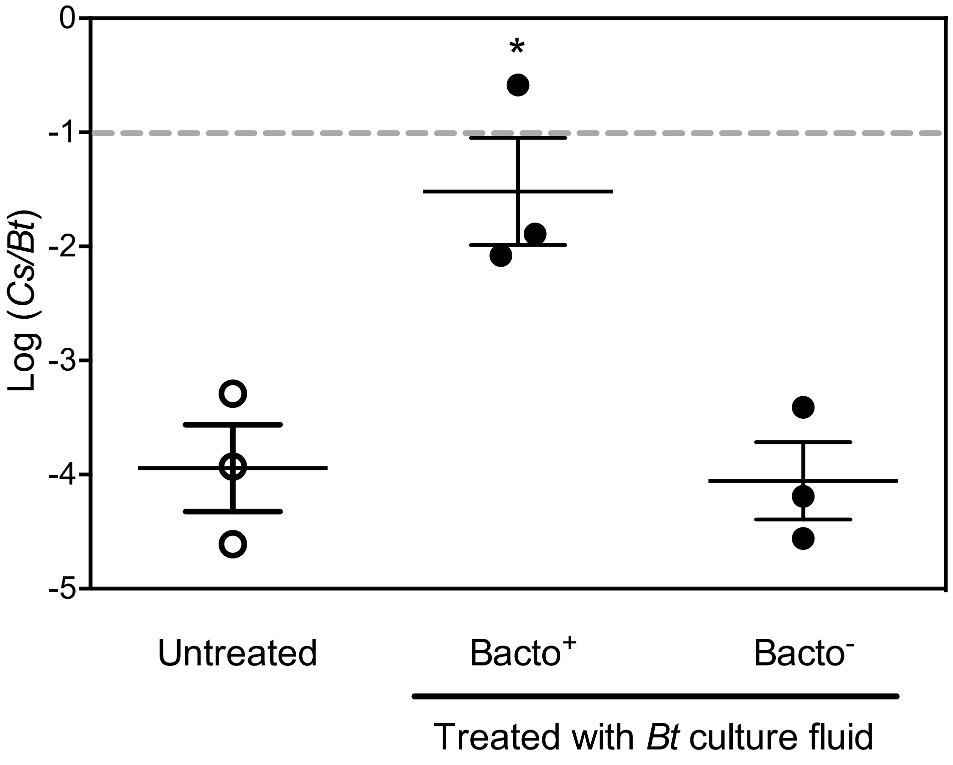

**Fig. S2. CdeR represses *cdeA* and *cdeB* transcription.** *cdeA* and *cdeB* transcripts in *C. substugae* wild type (WT, black bars) or a Δ*cdeR* mutant (white bars) stationary-phase cells (optical density of 600 nm [OD600] of 4). In all cases, results were normalized to the housekeeping gene encoding glyceraldehyde-3-phosphatase dehydrogenase (gapdh). The values represent the average of three independent experiments and the error bars represent the standard error of the mean. *, statistically significant by student's *t*-test ($p < 0.05$).

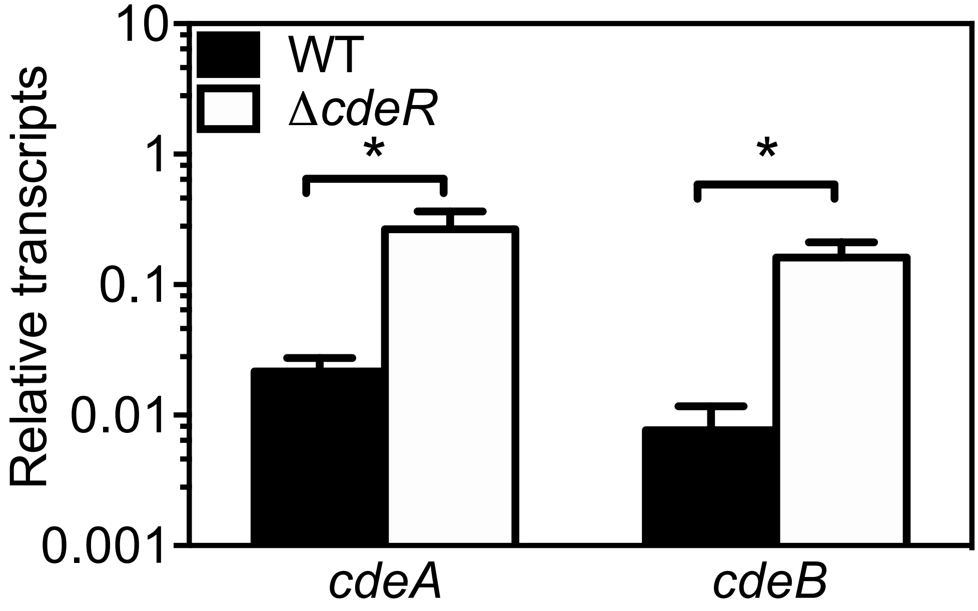

**Fig. S3. Ciprofloxacin and erythromycin do not significantly induce *C. subtsugae* tolerance.** Antibiotic minimum inhibitory concentration (MIC) was determined of cells treated for 6 h with antibiotic at a sublethal concentration (1/2 MIC from Table 1) (white bars) or identically treated cells with no antibiotic (black bars). The same antibiotic was used for both sublethal exposure and to assess MIC and is indicated by the X axis. Final MIC is shown as the average and standard error of 3 biological replicates for each test condition. For each antibiotic, there was no significant difference from untreated by *t*-test ($p$ > 0.2).

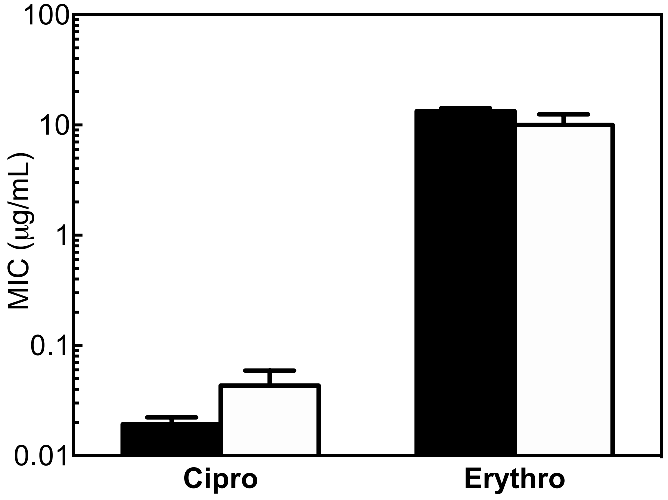

**Fig. S4. Bactobolin tolerance of *C. subtsugae* CV017 strains expressing CseR.** Antibiotic minimum inhibitory concentration (MIC) was determined of cells following 6 h incubation with bactobolin at a sub-MIC (1/2 MIC from Table 2) (white bars), or identically treated cells with no antibiotic (black bars). Final MIC is shown as the average and standard error of 4 biological replicates for each strain. For all strains, gentamicin antibiotic was included at every growth stage to maintain selection of the plasmid. *, statistical significance by students *t*-test ($p < 0.02$).

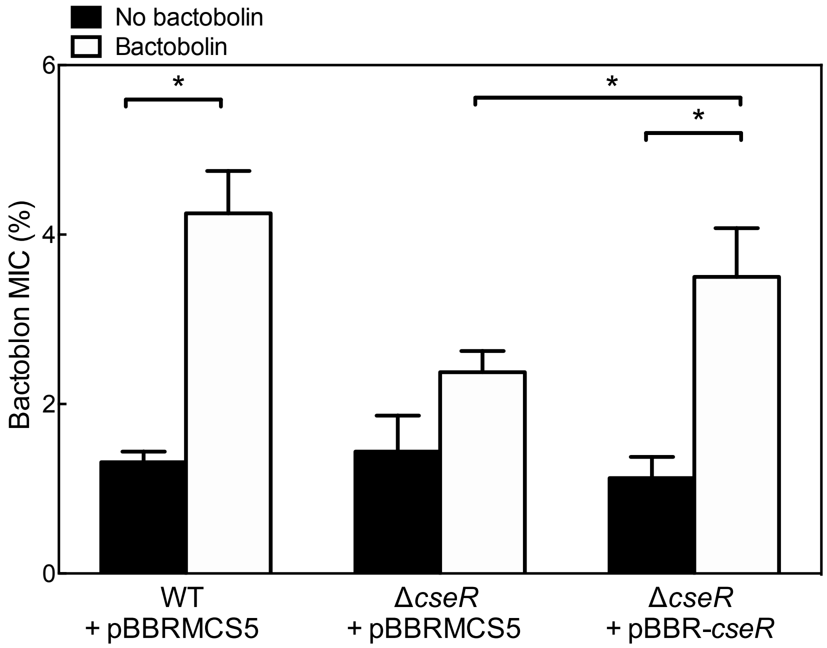

**Fig. S5. Phylogenetic tree of *Chromobacterium* species.** Tree is identical to that shown in Fig. 1, but here all species are shown. The scale indicates the number of substitutions per residue. Bootstrap values as the percentage of 1000 samples based on neighbor joining (NJ), maximum likelihood (ML) and Bayesian (Bay) methods are indicated for each node as defined in the figure legend. Nodes where all three methods showed values of >95% are indicated with a *. Red box indicates species encoding the *cseAB-oprN* genes. Red arrow, node where species with the *cseAB-oprN* gene cluster separate from other species.

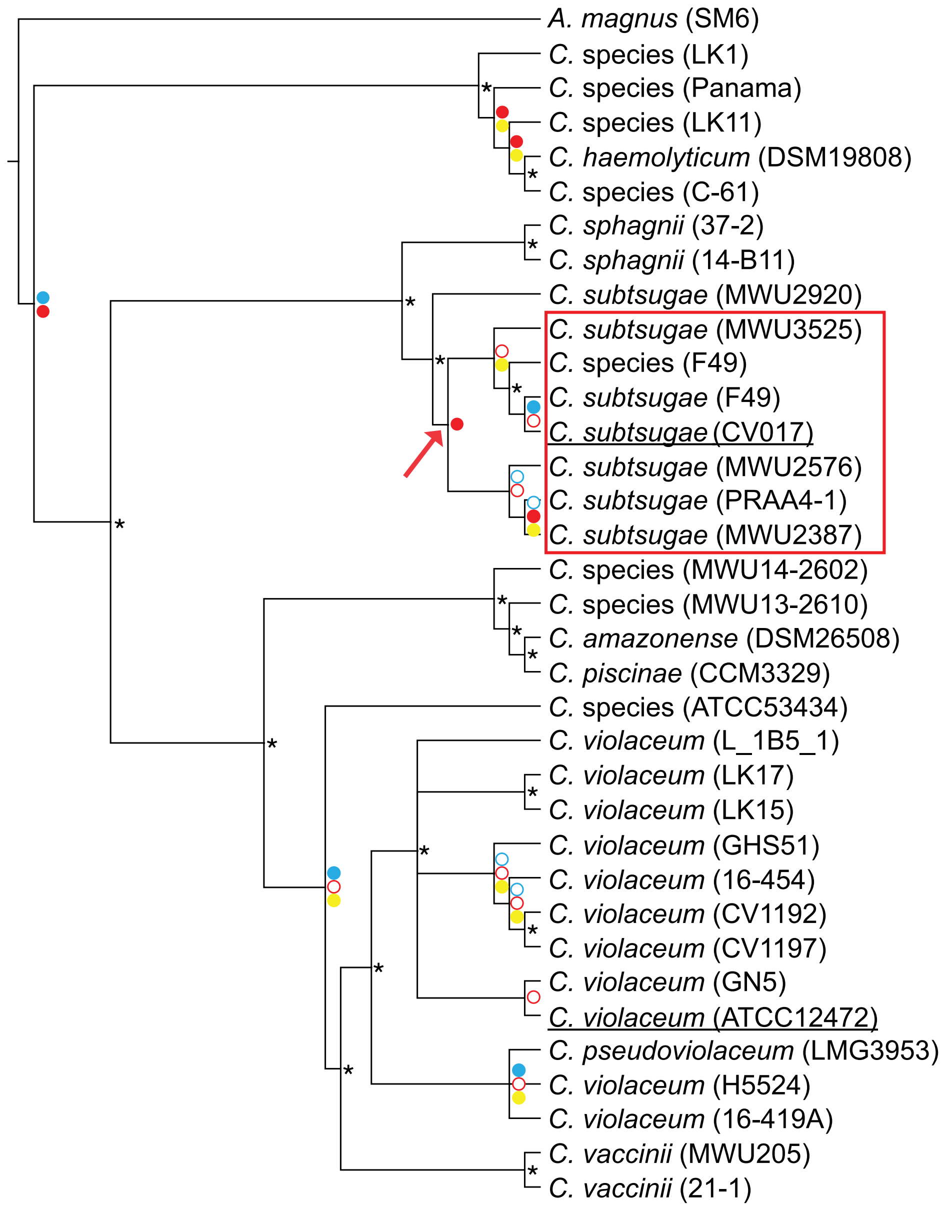

**Fig. S6. Phylogenetic tree of *Chromobacterium* cdeB *and* cseB *genes.*** The scale indicates the number of substitutions per residue. Bootstrap values as the percentage of 1000 samples based on neighbor joining (NJ), maximum likelihood (ML) and Bayesian (Bay) methods are indicated for each node as defined in the figure legend. Nodes where all three methods showed values of >95% are indicated with a *. Red box indicates species encoding the *cseAB-oprN* genes.

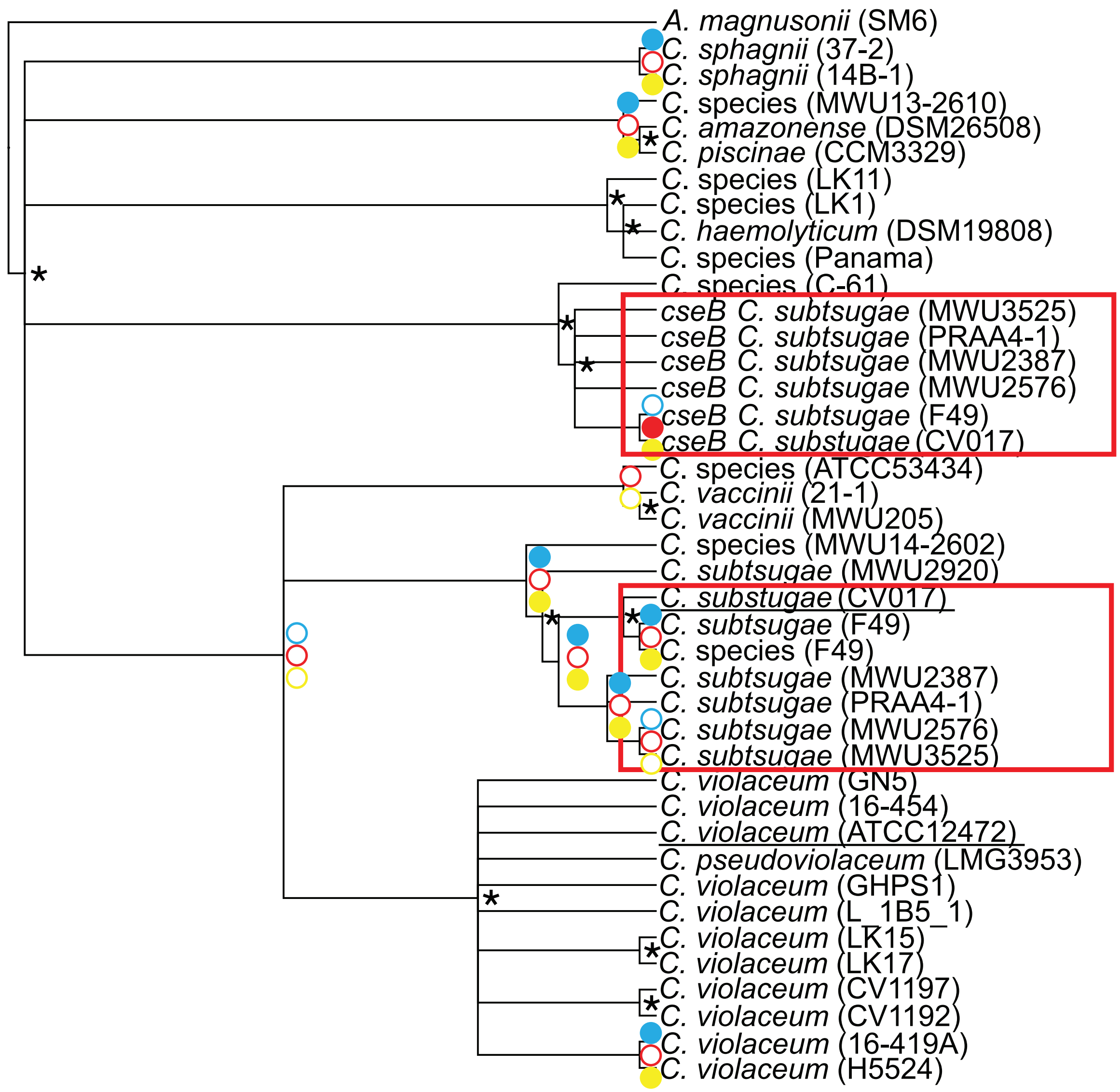
